# Estimating Effect Sizes and Expected Replication Probabilities from GWAS Summary Statistics

**DOI:** 10.1101/032474

**Authors:** Dominic Holland, Yunpeng Wang, Wesley K. Thompson, Andrew Schork, Chi-Hua Chen, Min-Tzu Lo, Aree Witoelar, Schizophrenia Working Group of the Psychiatric Genomics Consortium, Enhancing Neuro Imaging Genetics through Meta Analysis Consortium, Thomas Werge, Michael O’Donovan, Ole A. Andreassen, Anders M. Dale

## Abstract

Genome-wide Association Studies (GWAS) result in millions of summary statistics (“z-scores”) for single nucleotide polymorphism (SNP) associations with phenotypes. These rich datasets afford deep insights into the nature and extent of genetic contributions to complex phenotypes such as psychiatric disorders, which are understood to have substantial genetic components that arise from very large numbers of SNPs. The complexity of the datasets, however, poses a significant challenge to maximizing their utility. This is reflected in a need for better understanding the landscape of z-scores, as such knowledge would enhance causal SNP and gene discovery, help elucidate mechanistic pathways, and inform future study design. Here we present a parsimonious methodology for modeling effect sizes and replication probabilities that does not require raw genotype data, relying only on summary statistics from GWAS substudies, and a scheme allowing for direct empirical validation. We show that modeling z-scores as a mixture of Gaussians is conceptually appropriate, in particular taking into account ubiquitous non-null effects that are likely in the datasets due to weak linkage disequilibrium with causal SNPs. The four-parameter model allows for estimating the degree of polygenicity of the phenotype – the proportion of SNPs (after uniform pruning, so that large LD blocks are not over-represented) likely to be in strong LD with causal/mechanistically associated SNPs – and predicting the proportion of chip heritability explainable by genome-wide significant SNPs in future studies with larger sample sizes. We apply the model to recent GWAS of schizophrenia (N=82,315) and additionally, for purposes of illustration, putamen volume (N=12,596), with approximately 9.3 million SNP z-scores in both cases. We show that, over a broad range of z-scores and sample sizes, the model accurately predicts expectation estimates of true effect sizes and replication probabilities in multistage GWAS designs. We estimate the degree to which effect sizes are over-estimated when based on linear-regression association coefficients. We estimate the polygenicity of schizophrenia to be 0.037 and the putamen to be 0.001, while the respective sample sizes required to approach fully explaining the chip heritability are 10^6^ and 10^5^. The model can be extended to incorporate prior knowledge such as pleiotropy and SNP annotation. The current findings suggest that the model is applicable to a broad array of complex phenotypes and will enhance understanding of their genetic architectures.

## INTRODUCTION

Many complex traits and common phenotypes have a genetic component that arises from large numbers of genetic loci (Visscher et al., 2012). The total effect of the genetic component on phenotypic expression is often substantial, as indicated by measures of heritability (Witte et al., 2014; Tenesa and Haley, 2013) obtained from twin and family studies and genome-wide association studies (GWAS) for multiple phenotypes. For example, heritability of schizophrenia is estimated to be ~80% from twin studies (Sullivan et al., 2003), ~64% from family studies (Lichtenstein et al., 2009; Wray and Gottesman, 2012), with a lower bound of ~33% from recent GWAS (Ripke et al., 2013a). For any phenotype, GWAS provide a platform for uncovering the underlying genetic architecture, but this poses a substantial challenge, compounded by the complexity of the datasets: ~10^4^−10^5^ individuals with ~10^7^ genetic markers (single nucleotide polymorphisms, or SNPs) in various levels of correlation (linkage disequilibrium, or LD), ~10^6^ of which are estimated to be independent (Pe’er et al., 2008; Dudbridge and Gusnanto, 2008), with multiple possible roles for SNPs in mechanistic pathways.

Mathematical modeling is important for statistical genetics to capture, both broadly and in detail, the complexity of the datasets (So et al., 2010; Stahl et al., 2012; Schork, 2002). Indeed, with the numbers of markers much larger than numbers of individuals in GWAS, modeling assumptions are required so as to estimate parameters of interest and thereby obtain realistic descriptions of the numbers, distributions, and effect sizes of causal SNPs – and the considerably larger number of SNPs in strong LD with causal SNPs – which in turn can assist in causal SNP discovery and individual risk prediction, and inform mechanistic understanding of genetic effects in phenotypic expression.

Better understanding of the genetic architecture of complex traits will be facilitated both by many more individuals being genotyped, by fine-mapping, and by developing more advanced and realistic modeling techniques. Standard GWAS approaches, however, are designed for discovering a small number of common variants with large effects (i.e., low polygenicity), and so are not optimized for analyzing the large numbers of small effects in highly polygenic traits. Thus, there is a need for the development of analytical methods appropriate for the many phenotypes that have, or are expected to have, high polygenicity (Schork et al., 2013; Andreassen et al., 2013a,b).

Recently, methods have been developed to explore the combined contributions of many low-penetrance effects that do not reach genome-wide significance at current sample sizes. These include polygenic risk score profiling (Purcell et al., 2009; Schizophrenia Working Group of the Psychiatric Genomics Consortium, 2014); mixed linear modeling to estimate the genetic variance in unrelated individuals, where the distribution of effect sizes is modeled as a single normal (Yang et al., 2010; Lee et al., 2011, 2012a); a related Bayesian hierarchical model where the z-scores (or summary statistics for SNP association with phenotype), given the effect sizes, are assumed to follow a single normal distribution (So et al., 2011); modeling the distribution of the estimated genetic variance of *known* discoveries for a trait as a mixture of exponentials distribution, analogous to a scale mixture of normals distribution for the regression coefficients (Park et al., 2011); and an analysis that combines this later work with polygenic risk score profiling and heritability estimates from GWAS (Chatterjee et al., 2013). Additionally, multivariate linear mixed models ahve been developed (Yang et al., 2011a; Zhou and Stephens, 2014; Speed and Balding, 2014). The focus here, however, is on standard univariate analysis, but the empirical method for estimating regression coefficients in replication samples is also applicable to those arising from multrivariate analysis.

Mixture densities (Efron, 2013), particularly mixtures of normals, have previously been used in various forms to estimate effect sizes from individual-level data (Meuwissen et al., 2001; Erbe et al., 2012; Goddard et al., 2009; Zhou et al., 2013). Here we expand on a version of this relatively simple model for the distribution of z-scores (Thompson et al., 2015), and apply it to genome-wide summary statistics for schizophrenia and also to putamen volume, which provides an illustrative contrast.

One of our main objectives is to model, in a descriptive and accurate yet parsimonious way, the distribution of genetic summary statistics of traits for which a significant portion of the genome is involved, and thus help illuminate the genetic architecture of polygenic traits. To test for accuracy, we present a methodology for non-parametric estimation of quantities of interest which can then be compared with model predictions. This includes obtaining realistic estimates of the true effect size given the observed z-score and minimal other information: sample size and the model parameters that capture the statistics of the distribution – essentially, estimates of the conditional expectancy of regression coefficients *β* given *z*. Of particular importance, we want accurately to predict replication probabilities in multistage GWAS, i.e., the probability that a given z-score in a discovery sample will pass a nominal p-value significance threshold in a replication sample (that might include the discovery sample as a subset), a quantity that has not hitherto been a focus of much research. This quantity requires knowing the full distribution of test-statistics. In line with the parsimony of the model, the parameters will be directly interpretable – for example, one gives an index of the polygenicity – and being able accurately to estimate them and their uncertainties is a central component in this study. These will be used, for example, in power calculations to predict the proportion of additive chip heritability (which in turn is the proportion of phenotypic variance explainable by additive genetic effects of common SNPs assayed by GWAS arrays) that is discoverable as a function of sample size. (Additive chip heritability arises from additive contributions to phenotypic variance from tagged SNPs; below we will interchangeably refer to proportion of chip heritability and proportion of tagged variance explained by genome-wide significant SNPs.) Other recently developed methods that enable estimating chip heritability and proportion of variance explained are LD Scoring (Bulik-Sullivan et al., 2015) and Additive Variance Explained and Number of Genetic Effects Method of Estimation (AVENGEME) (Palla and Dudbridge, 2015).

It is large effects that GWAS have discovered in recent years, yet for many phenotypes it appears that very large numbers of small effects remain unidentified (Yang et al., 2010; Ripke et al., 2013a,b; Sklar et al., 2011). Thus, it will be necessary to include large (or sparse) and small (or ubiquitous) effects, a breakdown that naturally can be captured in a mixture model for the distribution of SNP z-scores: one Gaussian for each. In these Gaussians, it is important to incorporate the allele SNP heterozygosity (variance of the allele count). Null and non-null effects distributed throughout the genome could be captured by using only a single Gaussian; a two-groups mixture of Gaussians distribution has been used for ubiquitous null and sparse non-null effects; modifying this slightly will additionally allow for ubiquitous non-null effects: dedicating one Gaussian to ubiquitous null and non-null (small) effects, and the other to sparse (large) non-null effects. Intuitively, sparse effects represent SNPs that are in strong LD with causal SNPs (or more generally, with SNPs that are mechanistically associated with the phenotype), while the ubiquitous non-null effects largely arise from weak LD with causal SNPs, and the null effects – effects that do not replicate – arise from environmental and error contributions.

Here, we develop a unified framework for power calculations, relying on only four parameters and their uncertainties, that enables prediction of effect sizes, replication probabilities, and fraction of chip heritability explained by genome-wide significant SNPs as a function of sample size. We show that model variations that do not take into account ubiquitous non-null effects do not provide a good match to actual data, and propose a minor modification with important implications for the discovery of SNPs affecting phenotypes. We apply the model to the Psychiatric Genomics Consortium (PGC) (Sullivan, 2010) schizophrenia sample: 35,476 cases and 46,839 controls across 52 separate substudies, with imputation of SNPs using the 1000 Genomes Project reference panel (1000 Genomes Project Consortium, 2010) for a total of approximately 9.3 million genotyped and imputed SNPs (Schizophrenia Working Group of the Psychiatric Genomics Consortium, 2014). We also apply the model to putamen volume using data from the Enhancing Neuro Imaging Genetics through Meta-Analysis (ENIGMA) consortium (Hibar et al., 2015), with 12,596 subjects and the same set of SNPs as for schizophrenia. For these two phenotypes, using nonparametric methods described herein, we directly compare empirical with model results for expected replication effect sizes, variances, and replication probabilities, as a function of sample size, and map out the estimated proportion of chip heritability explained by genome-wide significant SNPs as a function of sample size.

## METHODS

### Proposed Gaussian Mixture Model

Assuming a linear relationship between a quantitative phenotype and genotype (logistic relationship for case-control designs), a massively univariate, or marginal regression, approach with effective sample size *N* (see Supplementary Material for definition) shows that the z-score for a given SNP can be written as a sum of genetic effect, *δ*, and a remainder term, *∊*, encompassing environmental and error contributions, assumed to be independent of *δ*: *z* = *δ* + *∊*, where 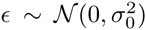, 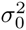 possibly being slightly different from 1 due to population substructure (Devlin and Roeder, 1999). In the context of the model, *δ* is the “true” effect size. Assuming Hardy-Weinberg equilibrium, for any particular SNP the effect size 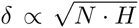, where *H* is the heterozygosity (allele count variance in the population), *H* = 2*p*(1 − *p*), *p* being the allele frequency for either of the two SNP alleles (see Supplementary Material for further details). Thus, the variance of *z* is 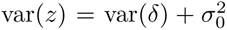, with var(*δ*) ∝ *N · H*. We introduce a four-parameter two-component Gaussian mixture model for the marginal distribution of z-scores assuming SNPs belong to a class of ubiquitous effects or to a class of sparse effects:

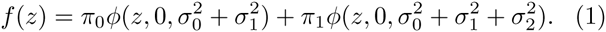

Here, π_0_ is the prior probability (after uniform pruning so that large LD blocks are not over-represented with respect to small LD blocks – see below) that a SNP is in the ubiquitous class (π_0_ ≈ 1) of “small” replicating effects (described by 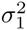); π_1_ = 1 − π_0_ is the prior probability that a SNP is in the sparse class of “large” replicating effects (described by 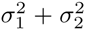), i.e. π_1_ is the fraction of independent SNPs characterized by the broader (sparse) normal probability distribution function (PDF), which we denote the index of polygenicity (π_1_ ≪ 1); 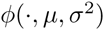 is the normal PDF with mean *μ* and variance *σ*^2^; all SNPs have a component that is a null effect, i.e., a non-replicating error/environmental contribution (associated with 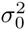); 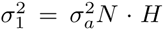 is the additional variance associated with non-null ubiquitous effects, and 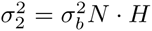 is the additional variance associated with sparse effects, with 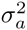 and 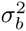 being the corresponding per-allele variances (assumed to be independent of allele frequency). The four parameters of the model then are π_1_, 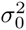, 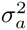, and 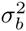. In the two-groups formalism, the non-null effect size is given by

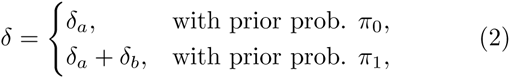
with *δ_a_* and *δ_b_* independent. The components of *δ* can be written as

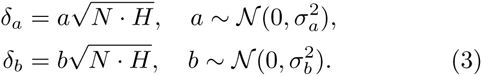

The more usual two-groups mixture model would have *σ_a_* ≡ 0, so that SNPs would be categorized as either truly null and ubiquitous, or non-null and sparse (Efron, 2013).

In the usual terminology applied to two-groups mixture models (Efron, 2013), the Bayesian local true discovery rate, *tdr*(*z*), is the probability that *z* corresponds to the *π*_1_ arm of Eq. 1, i.e., the posterior probability that a SNP with this z-score has a sparse effect:

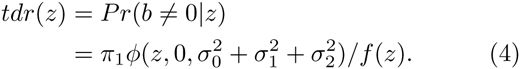

Then, from Eq. 1, it is easy to show that the posterior distribution of effect sizes *δ* given *z* is a weighted sum of sparse and ubiquitous contributions

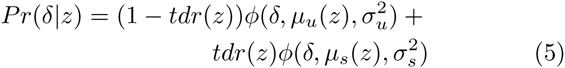
where,

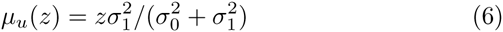

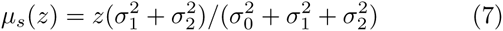

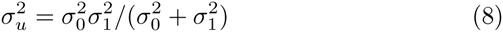

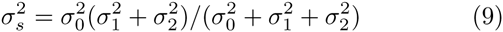
(see Supplementary Material for details). Note that from Eqs. 2, 3, and 5,

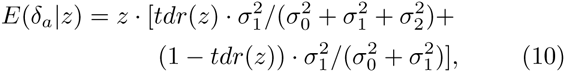

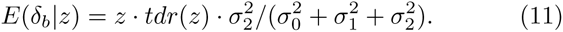
where *E* denotes expectation. Alternatively, based on Eq. 5, we can write *δ* = *δ_u_* + *δ_s_* for non-sparse and sparse effects, respectively, where

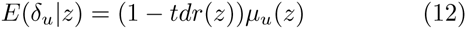

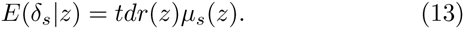

The objective, then, is to determine the empirical distribution of z-scores and the four model parameters that best characterize the data, assess the quality of the resulting model fit, and address its implications. Below we describe how the replication probability can be measured empirically. Of particular interest will be the accuracy of the model prediction of this quantity.

As there might be confuision about the terms bias and effect size, a note of clarification is in order. A true regression coefficient *β* for association between genotype and phenotype (see Supplementary Material) can be thought of as a true effect size. An estimate 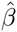 of this from data will then be an estimated effect size. Simple linear regression, the standard approach in GWAS, provides an unbiased estimate of effect sizes. That is, the marginal expectation of 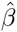 is 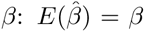. This is usually what is understood when the term “unbiased” is used for estimated quantities. However, the quantity of practical interest in association studies, because it is directly related to predicted z-scores in replication samples, is the conditional expectancy of the true effect size, given the estimate from GWAS: 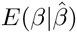. This quantity we call the adjusted effect size; our results below show that 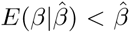. In other words, 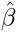 provides an inflated estimate of the true effect size. We show below that the adjusted effect size can be expressed as a function of 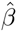, sample size *N*, heterozygosity *H*, and the four model parameters: 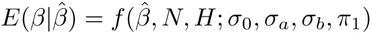. Since the Wald statistic (or z-score) corresponding to 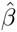 is simply a scaling of 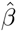, 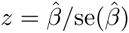, we will also refer to the component *δ* of *z* as the adjusted effect size, with context making it clear what is meant. (Note that “effect size” is also often used to denote the portion of phenotypic variance due to genotype, i.e., 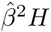 (Park et al., 2011)). We will show below that *δ* provides an unbiased estimate for the true effect size: given a discovery sample z-score *z_d_* for a particular SNP, then the expected value for the z-score in a replication sample is 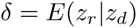.

### Empirical Estimation

We analyzed summary statistics (z-scores) from the PGC schizophrenia sample of 35,476 cases and 46,839 controls across 52 separate substudies, with 9,279,485 genotyped and imputed SNPs; restricting allele frequency to be greater than 0.005 reduced the number of SNPs by 2% (to 9083435). We randomly and repeatedly divide the data into complementary discovery and replication sets, and calculate the empirical expected z-score, and the expected square of the z-score, in the replication set, given a z-score in the discovery set. We also calculate empirical posterior estimates of the variance of the effect size in the replication set given a z-score in the discovery set, and the replication probability (defined below) for z-scores in the discovery set.

Note that while *π*_1_ and 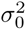 do not depend on sample size *N*, 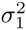 and 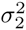 are proportional to *N*. Thus, to separate out these contributions and to examine the effects of sample size on posterior expectation values, the division of the data into complementary discovery and replication sets is not just a single division, e.g., split-half. Rather, we repeatedly and randomly divide the data with 10% of the effective sample in the discovery set (90% replication), then do the same in increments of 10%, through 90% of the effective sample in the discovery set (10% in replication). In the current work we do not use raw genotype data, but summary statistics from 52 studies with uneven distribution of effective sample sizes. So, for example, random draws are made from the 52 studies so that the studies selected for the discovery set will comprise approximately 10% of the total effective sample size, and so on for the other percentage breakdowns. The number of random draws per each percentage breakdown was 100, sufficient to provide smooth empirical a posteriori estimates of the quantities of interest. For each random draw, the SNPs were randomly pruned: for SNPs in LD, corresponding to correlation coefficient *r*^2^ ≥ 0.8, the SNP selected was randomly chosen – not necessarily the one with largest z-score. Using a fixed correlation coefficient allows for uniform pruning such that each LD block is treated equivalently (large block not over-represented); randomly selecting the representative SNP explicitly avoids the “winner’s curse” (Zöllner and Pritchard, 2007; Ghosh et al., 2008) in estimating discovery sample z-scores. Pruning at *r*^2^ ≥ 0.8 reduced the number of SNPs analyzed from ~ 9.3 to ~ 2.8 million, i.e., ~ 30% of the total.

Note, each iteration of the procedure produces an unbiased estimate of the posterior effect size means and variances, conditional on the discovery z-scores. The purpose of averaging across 100 random iterations is to smooth out the random differences present in each arbitrary partition of the sample into discovery and replication samples. Since each iteration is unbiased, the average across all iterations is again unbiased for the conditional posterior means and variances.

### Empirical Posterior Effect Sizes and Variances

For a given discovery-replication division of the data, z-scores from the discovery set were binned in 200 equally-spaced bins between *z_min_* = −6 and *z_max_* = 6. For each bin, the mean z-score for the corresponding SNPs was estimated from the replication set. For a given bin, denote the mean z-score in the discovery set as *z_d_*, and the corresponding mean z-score in the replication set as *z_r_*. Averaging these over the 100 repetitions (for a given percentage breakdown) provides an empirical estimate for the posterior expectation value of *z_r_* given *z_d_*, *E*(*z_r_*|*z_d_*). Note that the empirical *z_r_*, being a mean in the replication set, is a direct estimate of the effect size *δ_r_* in the replication set corresponding to *z_d_* in the discovery set: *E*(*z_r_* |*z_d_*) = *E*(*δ_r_* |*z_d_*). Similarly, empirical estimates for 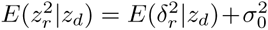 and 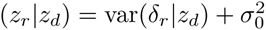 were calculated.

### Empirical Replication Probabilities

Given a discovery sample z-score *z_d_* for some SNP, the effect can be deemed to replicate if the corresponding z-score *z_r_* in the replication sample has the same sign as *z_d_*, and its p-value from a one-tailed test, based on the standard normal cumulative distribution, is less than a chosen threshold, say *p_t_* = 0.05, corresponding to – |*z_r_*| < *z_t_* = −1.645. For binned discovery sample z-scores, the empirical replication probability for a given bin is defined as the fraction of z-scores in the bin that replicate, i.e., have replication-sample p-value *p_r_* < *p_t_*. As before, averaging these over the 100 repetitions (for a given percentage breakdown) provides an empirical estimate of the replication probabilities, *R*(*z_d_*; *z_t_*), which we also denote *Pr*(*p_r_* < *p_t_*).

### Model Posterior Effect Sizes and Variances

For a discovery sample of effective sample size *N_d_* and a new replication sample of effective sample size *N_r_*, and noting that effect sizes are proportional to the square root of effective sample sizes, the posterior distribution for *z_r_* given *z_d_* is given by a modification of Eq. 5:

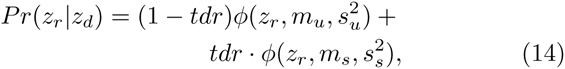
where

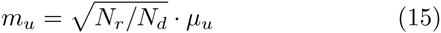

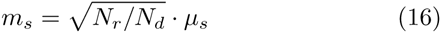

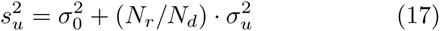

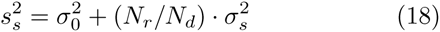
(the explicit dependence of *tdr*, *μ_u_*, and *μ*_s_ on *z_d_* and *N_d_*, and of *σ_u_* and *σ_s_* on *N_d_*, has been dropped to simplify the notation). Since *z* = *δ* + ∊, with *δ* and e assumed to be independent and *E*(*∊*) = 0, the expected effect size *δ_r_* in the replication sample, given a z-score *z_d_* in the discovery sample, can be read off from Eq. 14:

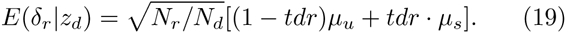

Additionally, from Eq. 14 and using standard properties of mixture distributions (see Supplementary Material), it also follows that

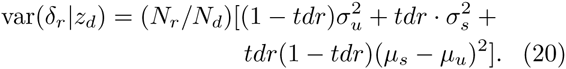

### Model Replication Probabilities

The replication rate *R*(*z_d_*; *z_t_*) for a given *z_d_* is the probability, in the complementary replication sample, of getting a z-score *z_r_* more significant than a chosen threshold, *z_t_* ≤ 0, which is simply the proportion of z-scores more significant than the threshold, which in turn is given by the cumulative distribution function (CDF) version of Eq. 14:

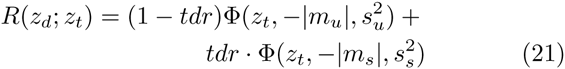
where Φ(*z_t_*, *m*, *s*^2^) is the normal CDF, corresponding to the normal PDF with mean *m* and variance *s*^2^, evaluated at *z_t_*.

### Parameter Estimation

The four model parameters were estimated by minimizing a convex cost function that was composed of a sum of two terms: the weighted sum of the squares of the differences between empirical estimates and model estimates of effect sizes (expected z-scores) and of the expected z-scores-squared (minimizing only with respect to the former does not allow for sufficient precision in determining the parameters). More specifically, denoting empirical replication z-scores as *z*, and model (predicted) posterior expectation values as *δ* and *η*, the cost function 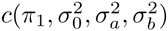 is

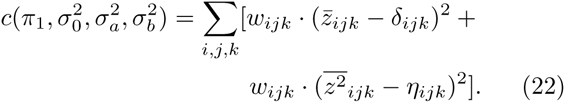

Here, the sum over *i* is over the 9 different percentage breakdowns of the dataset into complementary discovery-replication fractions; the sum over *j* is over the 100 repetitions for each breakdown; and the sum over *k* is over the 200 discovery sample z-score bins. For a given percentage breakdown *i* and repetition *j*, 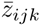 is the mean empirical replication z-score for discovery sample SNPs in bin *k* (note again that the mean replication z-score for a given bin provides a direct estimate of the effect size), and 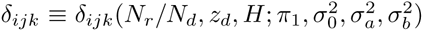 is the corresponding model prediction (only weakly dependent on repetition *j* through variation in heterozygosity *H* from repetition to repetition); 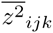 is the mean of the empirical replication z-score-squared for discovery sample SNPs in bin *k*, and *η_ijk_* is the corresponding model prediction. For given *i* and *j*, the weighting *w_ijk_* is the number of SNPs (z-scores) in the *k*-th bin. With *i* indexing the ratio *N_r_*/*N_d_* and *k* indexing *z_d_* (and dropping the explicit dependence of *δ_ijk_* and *η_ijk_* on *N_r_/Nd*, *H*, and the model parameters, so as to simplify the notation), 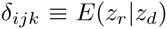 is given by Eq. 19 (*E*(*∊*) = 0), and 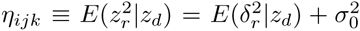 can be calculated from Eqs. 19 and 20, noting that

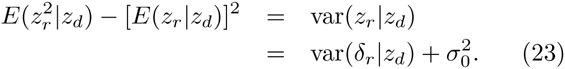

In particular, for a given sample of effective size *N*, the posterior expectation of the square of the “true” effect sizefor that sample is

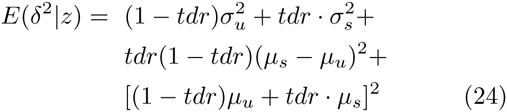
(note again that the explicit dependence of *tdr, μ_u_*, and *μ_s_ z* and *N*, and of *σ_u_* and *σ*_s_ on *N*, has been dropped to simplify the notation). Best-fit model parameters were determined by Nelder-Mead minimization of 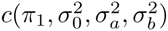, using the Matlab function `fminsearch()`.

### Proportion of Genetically-Determined Phenotypic Variance Explained, as a function of *N*

For a given sample size *N*, the expected proportion of the total additive genetic variance (approximately the proportion of chip heritability) explained by sparse effects, *S*(*N*; *z_t_*), for SNPs with – |*z*| < *z_t_* for some threshold *z_t_* < 0, or equivalently with p-value less than the corresponding threshold *p_t_*, can be estimated by simulating z-scores for all SNPs, whereby the unobservable null, ubiquitous, and sparse effects can explicitly be assigned to individual SNPs. From Eq. 1 and the implicit decomposition *z* = *δ* + *∊*, all of the simulation SNPs *i* (*i* = 1,…, ≃ 2.8 × 10^6^) are assigned an environmental/error component ei drawn from a normal distribution with mean zero and variance given by the estimated 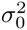. Then, a proportion *π*_i_ of the SNPs (indexed by *k*) – conceptually, SNPs that are in strong LD with causal SNPs – are assigned an additional component *δ*_c,_*_k_* drawn from a normal distribution with mean zero and variance 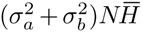, where 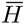 is the mean heterozygosity, so that the corresponding z-score for such a SNP *k* is *z_k_* = *S_c,k_* + *∊_k_*. (Though not necessary for the calculation of *S*, the remaining proportion, 1 – *π*_1_, of the SNPs can be assigned an additional (ubiquitous) component *δ_u,j_* drawn from a normal distribution with mean zero and variance 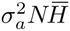, giving z-scores 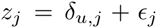. Note that the total effect size for a SNP in the *π*_1_ category can be decomposed into “ubiquitous” and “sparse” components: *δ_c,k_* = *δ_u,k_* + *δ_s,k_*.) The proportion of chip heritability explained by SNPs with sparse effects is given by the ratio

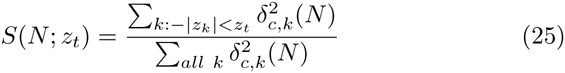
where 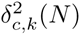 denotes the square of the “true” effect size component of the z-score for the *k*th SNP, emphasizing its dependence on *N* (see Supplementary Material). The numerator and denominator can be averaged over several repetitions for a smooth estimate of *S*(*z_t_*; *N*). Alternatively, given a set of z-scores {*z_k_*}, replace 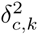 with the expectation 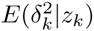 (see Eq. 24). The corresponding ratio in Eq. 25 should be accurate if the average effect of LD cancels between the numerator and denominator, which will always occur for *N* large enough so that *S* approaches 1. In any case, the effects of LD can increasingly be mitigated by higher levels of pruning.

### Multistage Design Combining Independent Discovery and Replication Datasets

It is common practice to estimate effect sizes in a multistage design where, at stage-1, candidate SNPs are selected in a discovery dataset using a liberal p-value (e.g., *p* = 10^−6^), and then reassessed at stage-2 in an independent replication data set, with the final assessment for significance being from the combined datasets (Lambert et al., 2013; Schizophrenia Psychiatric Genome-Wide Association Study (GWAS) Consortium, 2011; Ripke et al., 2013a). This strategy is usually employed when the independent replication dataset is not directly available to researchers, but where z-scores for candidate SNPs can be requested. The model presented here allows for predictions in this scenario. Specifically, let *N_d_* and *N_r_* be as before (sample sizes for independent discovery and replication datasets, respectively), and let *N_dr_* = *N_d_* + *N_r_* denote the sample size for the combined dataset. Define the inverse-variance weights 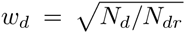 and 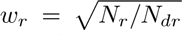. Then, given a discovery sample z-score *z_d_*, the posterior expectation for the z-score *z_dr_* in the combined dataset is

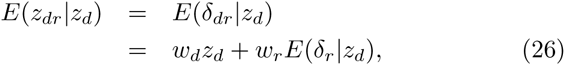
where *δ_dr_* is the effect size and *E*(*δ_r_* | *z_d_*) is given by Eq. 19.

The variance is simply

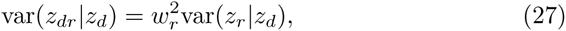
where 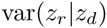 is given by Eqs. 20 and 23, thus allowing 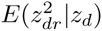 to be calculated. The replication probability 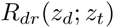 in the combined dataset for *z_d_* in the discovery dataset, which we can also write as *Pr*(*p_dr_* < *p_t_*) where *pdr* is the p-value for the SNP in the combined data set, is given by Eq. 21, only replacing *z_t_* on the right-hand side with 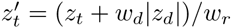:

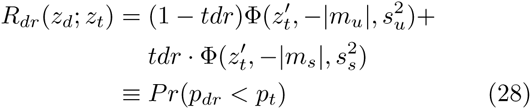
(see Supplementary Material for further details). Thus, for a SNP whose discovery sample z-score is *z_d_*, the probability of its z-score in the combined dataset reaching geome-wide significance is given by *R_dr_*(*z_d_*; *z_t_*), with *z_t_* = –5.33 (i.e., *Pr*(*p_dr_* < *p_t_*) with *p_t_* = 5 × 10^−8^).

## RESULTS

For a range of z-scores in the discovery sample between -6 and 6, Fig. 1 shows the empirical estimates (solid black curves) for schizophrenia of (A) expected effect sizes and (B) variances in the replication sample, and (C) the replication rate at *z_t_* = −1.64 (i.e., *pt* = 0.05), for split-half discovery and replication samples. It is significant that in (A), in a neighborhood of approximately ±3 around *z_d_* = 0 the expected effect size is non-zero: the black line has a positive slope. This implies that there exists ubiquitous non-null “small” effects (small z-scores have high probability densities, i.e., the corresponding SNPs are highly abundant). As the neighborhood extends further, “large” effects found in the discovery sample correspond to “large” effects in the replication sample.

**Figure 1:**
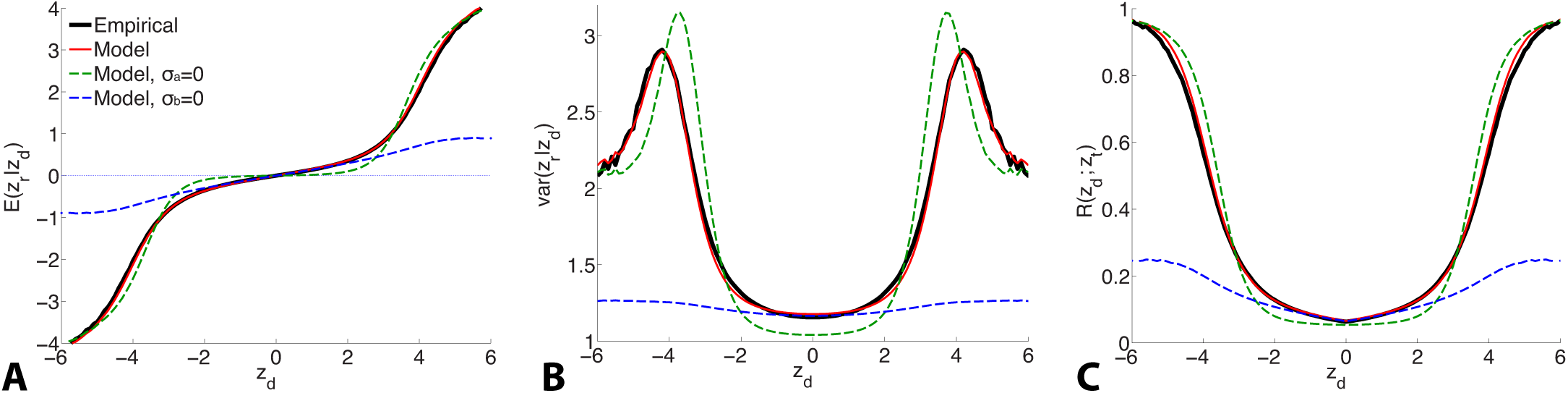
For schizophrenia, posterior estimate of (A) effect size and (B) variance; (C) estimate of replication probability for *z_t_ =* −1.64 (i.e., *p_t_ =* 0.05): empirical (black solid lines), current model (red solid lines), model with no ubiquitous effects (green dashed lines), and model with no sparse effects (blue dashed lines), for split-half discovery and replication data.

The parameter estimates (with 95% confidence intervals in square brackets) for schizophrenia were:

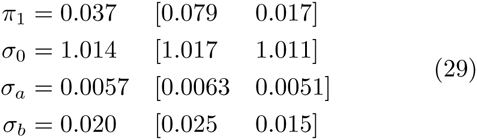
(the procedure for calculating the standard errors is described in the Supplementary Material). The model fit with these parameters is shown as the solid red curve in Fig. 1: there is an excellent fit to posterior effect size and variance, and to the replication rate. In contrast, if the model assumes no non-null ubiquitous effects (*σ_a_* = 0), then without changing the other parameters the fit corresponds to the green dashed line: in the ±3 neighborhood around *z_d_* = 0, small discovery effects do not replicate (i.e., they are null), while sparse “large” effects do replicate, in only approximate agreement with the empirical estimates. The alternative scenario assumes no sparse effects, *σ_b_* = 0: without altering the other parameters, the model fit is shown by the blue dashed curve in Fig. 1. In this case, the model provides a reasonable match with the empirical small effects. However, it completely fails to capture the posterior empirical effects for large *z_d_*. Corresponding curves for explicit model fits with parameters 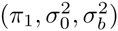 and 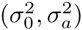, respectively, are shown in Supplementary Material Figure S1 (note that in the case of *σ_b_* =0 the model is degenetrate with respect to *π*_1_).

For putamen volume, the parameter estimates (with 95% confidence intervals) were:

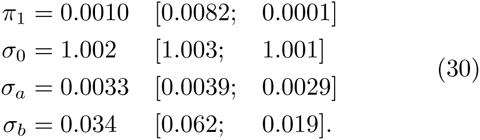

Thus, putamen volume is approximately 40-times less polygenic than schizophrenia.

Discovery and replication sample sizes have a pronounced influence on the empirical estimates of effect size, variance, and replication rate, but these dramatic changes across a wide range of z-scores are remarkably well matched by the model estimates (Supplementary Material, Fig. S3). Additionally, the empirical PDF for the z-scores is accurately reproduced by the model fit, Eq. 1, for z-scores divided up into five heterozygosity windows (Supplementary Material, Fig. S2), validating the basic model definition, Eq. 1.

Fig. 2 shows the posterior effect size components *δ_u_* (green curve) and *δ_s_* (red curve), given by Eqs. 12 and 13, for z-scores between -6 and 6, for an effective sample size *N* ≃ 34, 000. Also shown is the total effect size (black) and the posterior variance of the effect size (blue curve). The individual components can be seen to behave as expected: the green curve shows the ubiquitous non-null effects increasing away from the origin, reaching a peak, and falling to zero for large values of *z*; the red curve shows the sparse effects component essentially flat near the origin, indicating a lack of sparse effects in this neighborhood, and then monotonically increasing, beginning near where the ubiquitous effects start falling to zero. The peaks in the variance can be seen to arise from the regions where sparse effects are already prominent but the small effects have not yet died off. (Note that the variance is drawn on the same scale as the effect sizes.)

**Figure 2:**
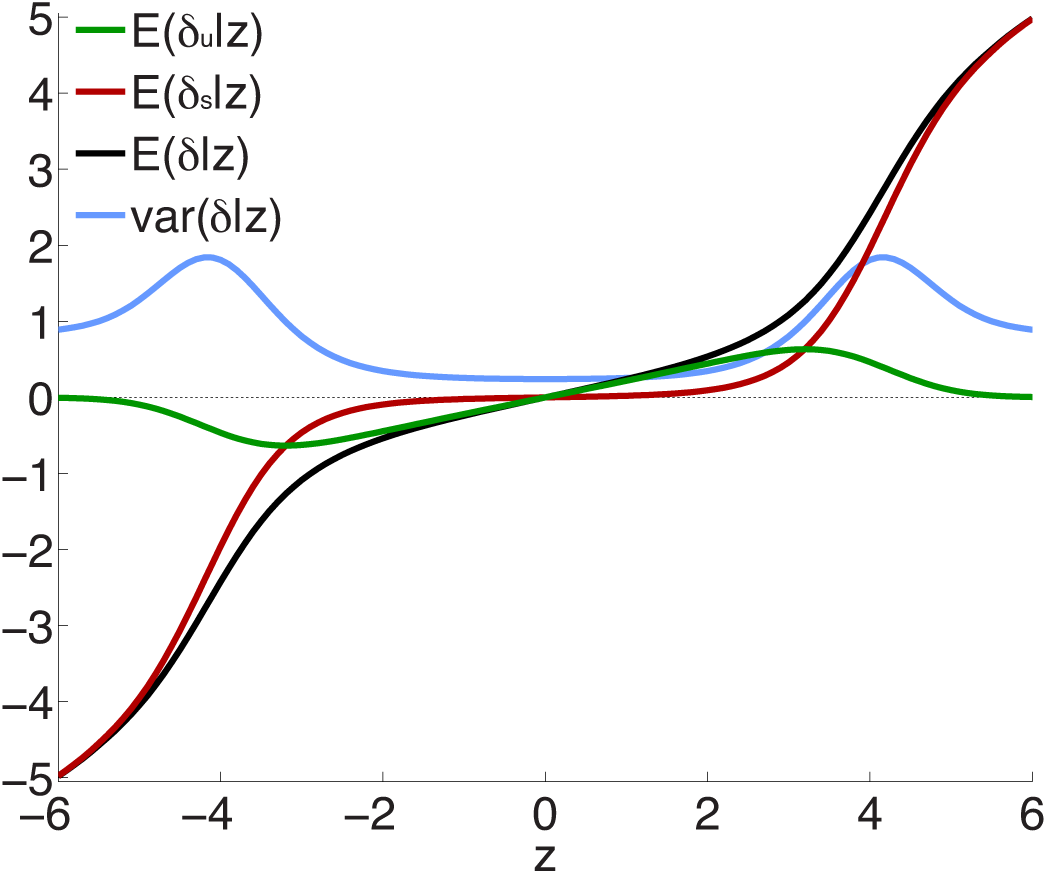
Posterior effect size and variance, calculated for effective sample size *N_d_* = *Nr* = *N* ≃ 34,000 – see Eqs. 12, 13, 20 and Supplementary Material. Note that sparse effects have a component that arises from ubiquitous effects. *z* = *δ* + *∊*, where *δ* = *δ_u_* + *δ_s_* and *E*(*∊*) = 0; *δ_u_* are ubiquitous effects, while *δ_s_* are additional contributions to total sparse effects.

To test the extent to which the small ubiquitous effects arose from LD with sparse large effects, in addition to light random pruning as before at *r*^2^ ≥ 0.8, we further restricted to SNPs whose total linkage disequilibrium (TLD), given by the sum of their LD *r*^2^’s with neighboring SNPs, was less than 15 (for reference, the median TLD was 54.5), reducing the number of SNPs to ≃ 1 million (10% of the total). The empirical effect sizes, for a sample breakdown of 50% discovery and 50% replication, are shown in Fig. 3 (the red plot). For comparison, also shown (in black) is the empirical effect size plot for random pruning at *r*^2^ ≥ 0.8 without restricting by TLD (same as in Fig. 1(A)). There is a pronounced diminution in the extent of ubiquitous (small) effects: the decreased slope near the origin suggests that a substantial portion of the ubiquitous effects arises due to LD with large effects, particularly large LD blocks (Yang et al., 2011b): if the distance between causal SNPs is comparable or smaller than typical LD block sizes, then “ubiquitous” effects are expected. Note that the shift from the black to the red curve in Fig. 3 is consonant with the shift from the black (ubiquitous and sparse effects) to the red (sparse effects only) curve in Fig. 2.

**Figure 3:**
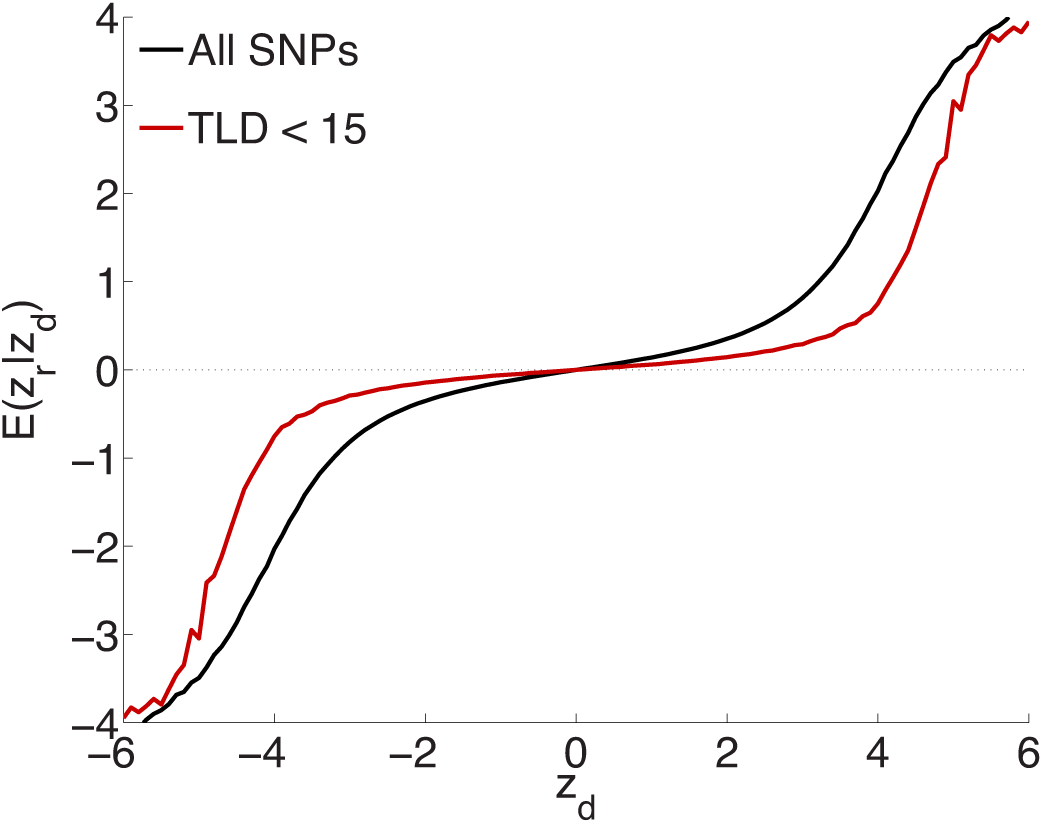
Randomly culling SNPs with LD *r^2^* ≥ 0.8 and further restricting to SNPs with total LD (TLD) less than 15 (approximately 1 million SNPs remaining) shows a diminution in the extent of ubiquitous effects (decreased slope near the origin for the red curve), consistent with an interpretation that the ubiquitous effects arise due to LD with causal SNPs. The black plot is for light random pruning at *r^2^ ≥* 0.8, shown in Fig. 1(A)

Fig. 4(A) shows quantile-quantile (QQ) plots for SNP p-values, comparing empirical and model fits for the full schizophrenia and putamen data sets, averaged over 100 repetitions with random pruning. For any threshold p-value, given as the ordinate in log_10_ units (to emphasize tail, or small p, behavior), the abscissa gives the proportion of SNPs whose p-values are at least as significant as the ordinate value; the dashed line at 45° corresponds to the null hypothesis where the distribution of SNP z-scores is assumed to follow a standard normal distribution (the threshold p-value then being synonymous with the proportion of SNPs exceeding that value). The model plots are given by Eq. 1, replacing the PDF with the CDF, and provide a remarkably good fit to the empirical plots. The earlier deviation of the schizophrenia plot, compared with the putamen plot, from the null line is due to the higher polygenicity of schizophrenia.

**Figure 4:**
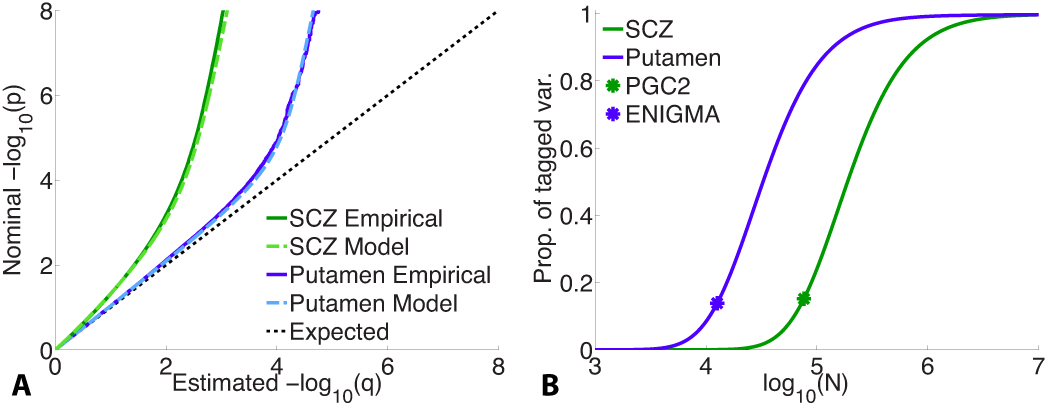
(A) Empirical and model QQ plots for putamen volume and schizophrenia. (B) Proportion of total additive genetic variance or chip heritability explained by sparse effects for all “tagged” SNPs with p-value less than the GWAS p-value threshold (*p_t_* = 5 × 10^−8^), as a function of effective sample size, for putamen volume and schizophrenia (the asterisks correspond to the current effective sample sizes for ENIGMA and PGS2). Of the total variance that is explained by sparse effects for all SNPs, the proportion explained by SNPs currently reaching the usual GWAS significance level is approximately 15% for both phenotypes.

Fig. 4(B) shows the projected proportion of tagged variance explained by SNPs reaching genome-wide significance (*p* ≤ 5 × 10^−8^) for schizophrenia and putamen volume, as a function of the total number *N* of subjects in the samples (assuming equal numbers of cases and controls for schizophrenia), as given by Eq. 25. The fraction of tagged variance explained by GWAS is expected to approximately equal the fraction of additive genetically-determined phenotypic variance, or narrow-sense heritability, explained. The blue asterisk indicates the sample size from current ENIGMA data (*N* = 12,596); the green asterisk, for schizophrenia, gives *N* = 76, 326, assuming the effective sample size from the current PGC2, *N_eff_* = 38, 163, arose from an equal number of cases and controls: *N*/2 – *N_cases_* = *N_controls_* = *N_eff_*. Thus we estimate that 15% of chip heritability for schizophrenia is currently explainable by genome-wide significant SNPs in PGC2; for these SNPs, the replication rate at *p_t_* = 0.05 for our split-half sample is 97% or higher. GWAS on approximately half a million each of cases and controls would need to be performed to explain all the chip heritability for schizophrenia. For putamen volume, 14% of chip heritability appears to be explainable by genome-wide significant SNPs given the current sample size in ENIGMA. In contrast to schizophrenia, however, approximately only 100,000 people would need to be assessed to fully explain chip heritability for putamen volume. The higher sample size requirements for schizophrenia are due partly to its higher polygenicity.

The per-allele contribution of a locus, with z-score *z*, to the phenotypic variance *v_a_* of the trait is usually estimated to be 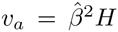, where 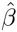 is the corresponding regression coefficient in the univariate setting (Park et al., 2010). This is proportional to *z*^2^/*N*, since 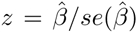. However, the “true” effect more correctly is related to the non-centrality parameter *δ* (*z* = *δ* + *∊*), so that the per-allele contribution to phenotypic variance is proportional to *E*(*δ*^2^|*z*)/*N*, not *z*^2^/*N*. Fig. 5(A) plots *E*(*δ*^2^|*z*) versus *z*^2^ (or scaled 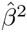) for three illustrative total sample sizes. For an independent sample, the degree by which *v_a_* is an overestimate when based on *z*^2^ instead of *E*(*δ*^2^|*z*) is given by the ratio of the height on the black dotted line to the height on the appropriate curve, at *z*^2^. For example, for schizophrenia with a total sample size of *N*=50,000 and a z-score on the threshold of genome-wide significance (*δ* ≃ ±5.33), *v_a_* will be over-estimated by a factor of 2.2 (the height of the black dot relative to the blue dot).

**Figure 5:**
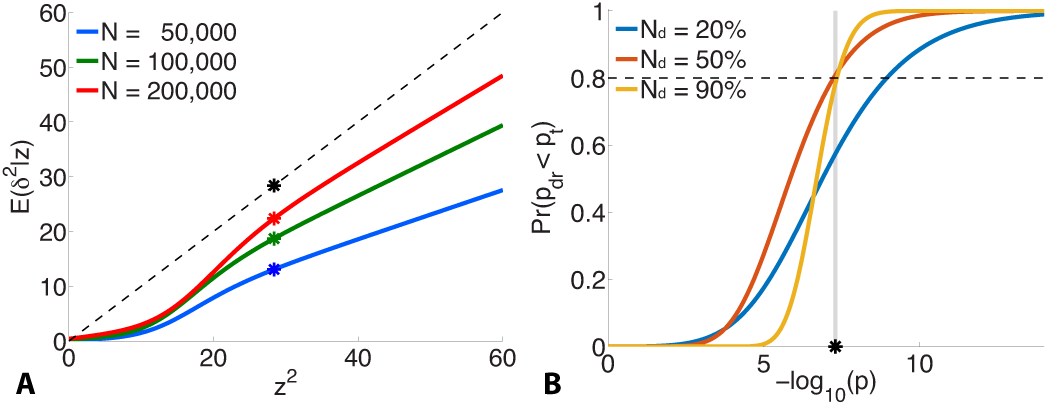
For schizophrenia, (A) posterior estimates of effect-size-squared, as given by Eq. 24, versus *z*^2^ for three total sample sizes. When assuming that the phenotypic variance explained by a SNP is given by 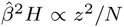, the degree to which this is an over-estimate is indicated by the ratio of the height of the black dashed line (the assumption *δ*^2^ = *z*^2^) to the height of the corresponding point on the curve for a given sample size. The asterisks correspond to the threshold significant z-score. (B) For a multistage GWAS, where discovery is from a subset (20%, 50%, 90%) of the total PGC2 sample, the curves give the probability of a SNP with p-value *p* in the discovery sample passing genome-wide significance (*pd_r_ < pt* = 5 × 10^−8^) in the combined (total) data set, Eq. 28. The vertical grey line is at *p* = *p_t._*

Fig. 5(B) shows the probability, given by Eq. 28, of reaching genome-wide significance in a combined discovery and replication dataset of total sample size corresponding to PGC2 (N = 76, 326), where the discovery sample size *N_d_* is 20%, 50%, or 90% of the total, as a function of p-value in the discovery dataset. For example, in discovery samples equal to 50% or 90% of the total, to have a probability of reaching genome-wide signifiance in the combined dataset approximately equal to 0.8 would require having reached genome-wide significance in the discovery dataset; for a discovery sample only 20% of the total, the same probability of replicating requires the discovery sample p-value to be at least *p* = 10^−9^, i.e., *more* significant than the genome-wide threshold.

## DISCUSSION

Here we present a simple modeling framework, a scale-mixture of Gaussians that is a modification of previously published methodologies (Efron, 2013; Meuwissen et al., 2001; Erbe et al., 2012; Goddard et al., 2009; Zhou et al., 2013), for assessing the contributions of ubiquitous and sparse effects to quantities of interest in GWAS. Additionally, we present a procedure for testing the model empirically. The model has great utility in that it allows for the prediction of the replication probability and expected effect size for each SNP, given the discovery and replication sample sizes, the four model parameters, and the discovery sample z-score of the SNP. Using nonparametric methods applied to large schizophrenia and subcortical brain structure volumes, we show how the empirical replication probabilities and effect sizes can be calculated directly from the data, and demonstrate that the model results are in excellent agreement with these – across a wide range of z-scores and sample sizes, and for multistage GWAS designs, whether splitting the available sub-studies into randomized disjoint sets for discovery and replication, or combining the discovery dataset as a subset in replication.

The parameter *π*_1_ in the model gives the prior probability that SNPs belong in the category of sparse effects, i.e., those likely to have a large and significant association with the phenotype. *π*_1_ thus provides a way of estimationg the proportion of “causal” SNPs. An alternative method for doing this is Approximate Bayesian Polygenic Analysis (ABPA) (Stahl et al., 2012). With 1000 Genomes SNP imputation for a total of approximately 9.3 million SNPs in the studies, all possible causal common SNPs are likely to be represented, either directly or through strong LD. These SNPs will have the largest expected association summary statistics; the remaining SNPs will either show weaker effects through attenuated LD with the causal SNPs, or have null effects. To a first approximation, a plausible interpretation is that SNPs in the *π*_1_ category are likely to be dominated by SNPs in strong LD with causal SNPs. Our estimate of *π*_1_ ≃ 0.037 for the PGC2 schizophrenia GWAS suggests that this condition is highly polygenic: about 3.7% of SNPs are potentially significantly associated with the phenotype. It should be noted, however, that variations at different loci exhibiting LD may have independent effects on a phenotype (Malo et al., 2008).

Recently, a schizophrenia GWAS study, using approximately half of the subjects and all of the SNPs employed here, reported an estimate of “SNP heritability” (a lower bound of narrow sense heritability since only variation due to SNP association can be determined; copy-number variants and rare variants, for example, are not included) *h*^2^ = 33% on the liability scale, adjusted for case-control ascertainment (Ripke et al., 2013a; Lee et al., 2012b). It should be noted that an implicit assumption in the method used for estimating *h*^2^ is that the distribution of effect sizes is given by a single Gaussian (no explicit sparse-effects component – see Figure 1(A): the blue dashed curve for *σ_b_* = 0), whereas we have shown here that it is more appropriate to consider the effect sizes being described by a Gaussian mixture distribution, with non-null ubiquitous and sparse components. It is not clear how this affects the result for *h*^2^. A follow-up study, using all of the subjects and SNPs employed here, reported that 3.4% of variaiton on the liability scale to schizophrenia, adjusted for case-control ascertainment, is explained by genome-wide significant loci (Schizophrenia Working Group of the Psychiatric Genomics Consortium, 2014). Therefore, approximately 10% (3.4/33) of SNP heritability (or chip heritability) for schizophrenia was estimated to be captured by genome-wide significant SNPs, a result that comports with our estimate of 15%. For the putamen, a recent report (Hibar et al., 2015) using all of the subjects and SNPs employed here, and a different method for estimating SNP heritability than was used for schizophrenia (So et al., 2011), found that approximately 10% of variance in putamen volume was due to all common variants, while 1.09% was attributable to genome-wide significant SNPs. Thus, again, approximately 10% of SNP heritability for putamen volume was estimated to be captured by genome-wide significant SNPs, a result in broad agreement with our estimate of 14%.

We presented a four-parameter two-groups mixture of normals parametric model for the distribution of GWAS summary statistics. Additionally, we presented an empirical scheme for estimating replication z-scores, their variances, and replication rates. The model parameters are then estimated by minimizing a cost function that depends on the differences in model and empirical estimates of effect sizes and expected z-scores-squared. Applying the model and empirical scheme to recent large GWAS of schizophrenia and putamen volume, we showed that effect sizes, along with variances and replication rates, are accurately described by the simple model over a wide range of z-scores, and over a wide range of discovery and replication sample sizes when the relationships can change dramatically. We further showed how the model can be used to estimate the fraction of additive SNP heritability (an estimate of the fraction of chip heritability) explainable by genome-wide significant SNPs in a massive univariate setting, as a function of sample size. The model enables estimation of an index for the degree of polygenicity of a phenotype, because structurally it allows for separate distributions of the large number of ubiquitous (small) effects and the relatively small number of sparse (large) effects, postulating that these different classes of effects will be distributed differently. We showed that ignoring the contribution of ubiquitous non-null effects severely degrades the accuracy of the mode. Given the model parameters, for any SNP with a p-value less than some threshold – indicating a potential candidate for true association with the phenotype – an accurate estimate of its probability of replicating can be calculated for replication samples of different effective numbers of subjects. It should be noted, however, that to have reasonable probability of replicating in a multistage GWAS, it might be necessary that candidate SNPs in the discovery sample substantially *exceed* genome-wide significance. The model can be improved and extended by letting the model coefficients depend on SNP heterozygosity, and by incorporating additional information like SNP functional annotation category (Schork et al., 2013). We showed that schizophrenia is highly polygenic (~3.7% of SNPs are significantly associated). We estimate that at approximately half a million each of cases and controls genotyped, all SNPs contributing to narrow-sense heritability would have reached genome-wide significance. Putamen volume is approximately 40-times less polygenic than schizophrenia, and only of order one hundred thousand people need to be genotyped to capture all chip heritability with genome-wide significant SNPs.

Deep sequencing of large samples can be expected to advance our understanding of the genetics of complex phenotypes, but it seems more cost-efficient to start with uncovering more risk genes from existing GWAS data by improving analytical tools, as much remains to be explored in the complex landscape where large numbers of SNPs in GWAS effect phenotype. In particular, there is a need for better understanding the distribution of effect sizes in GWAS (Schork et al., 2013; Andreassen et al., 2013a,b). The univariate mixture model we have presented and validated empirically here captures the distribution of SNP effect sizes, and thus much of the genetic architecture of complex phenotypes. It can be used for estimating polygenicity and informing power calculations, in particular for providing estimates of probabilities of reaching genome-wide significance in multistage GWAS, and the proportion of SNP heritability explainable in future larger studies. The model can be extended by incorporating additional prior information, such as SNP annotation category. Combined with larger sample sizes, this statistical methodology may facilitate improved detection of smaller effect sizes and enhance the ability of GWAS in accurate risk prediction, and to inform human physiology and disease etiology.

## Acknowledgments

The authors wish to thank the participants of the studies for their contribution, as well as all researchers in the Schizophrenia Working Group of the Psychiatric Genomics Consortium and in the Enhancing Neuro Imaging Genetics through Meta Analysis Consortium who contributed with GWAS data. A full list of these researchers and their affiliations is provided in the Supplementary Material.

## Funding

The study was financially supported by NIH, NIH GM104400-01A, the Research Council of Norway (#213837, #223273), the South-East Norway Regional Health Authority (#2013-123) and KG Jebsen Foundation.

## References

1000 Genomes Project Consortium, 2010. A map of human genome variation from population-scale sequencing. Nature 467 (7319), 1061–1073.

Andreassen, O. A., Djurovic, S., Thompson, W. K., Schork, A. J., Kendler, K. S., O’Donovan, M. C., Rujescu, D., Werge, T., van de Bunt, M., Morris, A. P., et al., 2013a. Improved detection of common variants associated with schizophrenia by leveraging pleiotropy with cardiovascular-disease risk factors. The American Journal of Human Genetics 92 (2), 197–209.

Andreassen, O. A., Thompson, W. K., Schork, A. J., Ripke, S., Mattingsdal, M., Kelsoe, J. R., Kendler, K. S., O’Donovan, M. C., Rujescu, D., Werge, T., et al., 2013b. Improved detection of common variants associated with schizophrenia and bipolar disorder using pleiotropy-informed conditional false discovery rate. PLoS genetics 9 (4), e1003455.

Bulik-Sullivan, B. K., Loh, P.-R., Finucane, H. K., Ripke, S., Yang, J., Patterson, N., Daly, M. J., Price, A. L., Neale, B. M., of the Psychiatric Genomics Consortium, S. W. G., et al., 2015. Ld score regression distinguishes confounding from polygenicity in genome-wide association studies. Nature genetics 47 (3), 291–295.

Chatterjee, N., Wheeler, B., Sampson, J., Hartge, P., Chanock, S. J., Park, J.-H., 2013. Projecting the performance of risk prediction based on polygenic analyses of genome-wide association studies. Nature genetics 45 (4), 400–405.

Devlin, B., Roeder, K., Dec 1999. Genomic control for association studies. Biometrics 55 (4), 997–1004.

Dudbridge, F., Gusnanto, A., 2008. Estimation of significance thresholds for genomewide association scans. Genetic epidemiology 32 (3), 227–234.

Efron, B., 2013. Large-scale inference: empirical Bayes methods for estimation, testing, and prediction. Cambridge University Press, Cambridge, UK New York.

Erbe, M., Hayes, B.,Matukumalli, L., Goswami, S., Bowman, P., Reich, C., Mason, B., Goddard, M., 2012. Improving accuracy of genomic predictions within and between dairy cattle breeds with imputed high-density single nucleotide polymorphism panels. Journal of dairy science 95 (7), 4114–4129.

Ghosh, A., Zou, F., Wright, F. A., 2008. Estimating odds ratios in genome scans: an approximate conditional likelihood approach. The American Journal of Human Genetics 82 (5), 1064–1074.

Goddard, M. E., Wray, N. R., Verbyla, K., Visscher, P. M., et al., 2009. Estimating effects and making predictions from genome-wide marker data.Statistical Science 24 (4), 517–529.

Hibar, D. P., Stein, J. L., Renteria, M. E., Arias-Vasquez, A., Desrivières, S., Jahanshad, N., Toro, R., Wittfeld, K., Abramovic, L., Andersson, M., et al., 2015. Common genetic variants influence human subcortical brain structures. Nature.

Lambert, J.-C., Ibrahim-Verbaas, C. A., Harold, D., Naj, A. C., Sims, R., Bellenguez, C., Jun, G., DeStefano, A. L., Bis, J. C., Beecham, G. W., et al., 2013. Meta-analysis of 74,046 individuals identifies 11 new susceptibility loci for alzheimer’s disease. Nature genetics 45 (12), 1452–1458.

Lee, S. H., DeCandia, T. R., Ripke, S., Yang, J., Sullivan, P. F., Goddard, M. E., Keller, M. C., Visscher, P. M., Wray, N. R., Mar 2012a. Estimating the proportion of variation in susceptibility to schizophrenia captured by common SNPs. Nat. Genet. 44 (3), 247–250.

Lee, S. H., Goddard, M. E., Wray, N. R., Visscher, P. M., 2012b. A better coefficient of determination for genetic profile analysis. Genetic epidemiology 36 (3), 214–224.

Lee, S. H., Wray, N. R., Goddard, M. E., Visscher, P. M., 2011. Estimating missing heritability for disease from genome-wide association studies. The American Journal of Human Genetics 88 (3), 294–305.

Lichtenstein, P., Yip, B. H., Björk, C., Pawitan, Y., Cannon, T.D., Sullivan, P. F., Hultman, C. M., 2009. Common genetic influences for schizophrenia and bipolar disorder: a population-based study of 2 million nuclear families. Lancet 373 (9659).

Malo, N., Libiger, O., Schork, N. J., 2008. Accommodating linkage disequilibrium in genetic-association analyses via ridge regression. The American Journal of Human Genetics 82 (2), 375–385.

Meuwissen, T. H. E., Hayes, B. J., Goddard, M. E., 2001. Prediction of total genetic value using genome-wide dense marker maps. Genetics 157 (4), 1819–1829.

Palla, L., Dudbridge, F., 2015. A fast method that uses polygenic scores to estimate the variance explained by genome-wide marker panels and the proportion of variants affecting a trait. The American Journal of Human Genetics 97 (2), 250–259.

Park, J. H., Gail, M. H., Weinberg, C. R., Carroll, R. J., Chung, C. C., Wang, Z., Chanock, S. J., Fraumeni, J. F., Chatterjee, N., Nov 2011. Distribution of allele frequencies and effect sizes and their interrelationships for common genetic susceptibility variants. Proc. Natl. Acad. Sci. U.S.A. 108 (44), 18026–18031.

Park, J.-H., Wacholder, S., Gail, M. H., Peters, U., Jacobs, K. B., Chanock, S. J., Chatterjee, N., 2010. Estimation of effect size distribution from genome-wide association studies and implications for future discoveries. Nature genetics 42 (7), 570–575.

Pe’er, I., Yelensky, R., Altshuler, D., Daly, M. J., 2008. Estimation of the multiple testing burden for genomewide association studies of nearly all common variants. Genetic epidemiology 32 (4), 381–385.

Purcell, S. M., Wray, N. R., Stone, J. L., Visscher, P. M., O’Donovan, M. C., Sullivan, P. F., Sklar, P., Purcell, S. M., Stone, J. L., Sullivan, P. F., et al., 2009. Common polygenic variation contributes to risk of schizophrenia and bipolar disorder. Nature 460 (7256), 748–752.

Ripke, S., O’Dushlaine, C., Chambert, K., Moran, J. L., Kähler, A. K., Akterin, S., Bergen, S. E., Collins, A. L., Crowley, J. J., Fromer, M., et al., 2013a. Genome-wide association analysis identifies 13 new risk loci for schizophrenia. Nature genetics 45 (10), 1150–1159.

Ripke, S., Wray, N. R., Lewis, C. M., Hamilton, S. P., Weissman, M. M., Breen, G., Byrne, E. M., Blackwood, D. H., Boomsma, D. I., Cichon, S., et al., 2013b. A mega-analysis of genome-wide association studies for major depressive disorder. Molecular psychiatry 18 (4), 497–511.

Schizophrenia Psychiatric Genome-Wide Association Study (GWAS) Consortium, 2011. Genome-wide association study identifies five new schizophrenia loci. Nature genetics 43 (10), 969–976.

Schizophrenia Working Group of the Psychiatric Genomics Consortium, Jul 2014. Biological insights from 108 schizophrenia-associated genetic loci. Nature 511 (7510), 421–427.

Schork, A. J., Thompson, W. K., Pham, P., Torkamani, A., Roddey, J. C., Sullivan, P. F., Kelsoe, J. R., O’Donovan, M. C., Furberg, H., Schork, N. J., et al., 2013. All snps are not created equal: genome-wide association studies reveal a consistent pattern of enrichment among functionally annotated snps. PLoS genetics 9 (4), e1003449.

Schork, N. J., 2002. Power calculations for genetic association studies using estimated probability distributions. The American Journal of Human Genetics 70 (6), 1480–1489.

Sklar, P., Ripke, S., Scott, L. J., Andreassen, O. A., Cichon, S., Craddock, N., Edenberg, H. J., Nurnberger, J. I., Rietschel, M., Blackwood, D., et al., 2011. Large-scale genome-wide association analysis of bipolar disorder identifies a new susceptibility locus near odz4. Nature genetics 43 (10), 977.

So, H.-C., Li, M., Sham, P. C., 2011. Uncovering the total heritability explained by all true susceptibility variants in a genome-wide association study. Genetic epidemiology 35 (6), 447–456.

So, H.-C., Yip, B., Sham, P. C., 2010. Estimating the total number of susceptibility variants underlying complex diseases from genome-wide association studies. PLoS One 5 (11), e13898-e13898.

Speed, D., Balding, D. J., 2014. Multiblup: improved snp-based prediction for complex traits. Genome research 24 (9), 1550–1557.

Stahl, E. A., Wegmann, D., Trynka, G., Gutierrez-Achury, J., Do, R., Voight, B. F., Kraft, P., Chen, R., Kallberg, H. J., Kurreeman, F. A., et al., 2012. Bayesian inference analyses of the polygenic architecture of rheumatoid arthritis. Nature genetics 44 (5), 483–489.

Sullivan, P. F., Oct 2010. The psychiatric GWAS consortium: big science comes to psychiatry. Neuron 68 (2), 182–186.

Sullivan, P. F., Kendler, K. S., Neale, M. C., Dec 2003. Schizophrenia as a complex trait: evidence from a meta-analysis of twin studies. Arch. Gen. Psychiatry 60 (12), 1187–1192.

Tenesa, A., Haley, C. S., 2013. The heritability of human disease: estimation, uses and abuses. Nature Reviews Genetics 14 (2), 139–149.

Thompson, W. K., Wang, Y., Schork, A., Zuber, V., Andreassen, O. A., Dale, A. M., Holland, D., Shujing, X., 2015. An empirical bayes method for estimating the distribution of effects in genome-wide association studies. PLoS Genetics [in press].

Visscher, P. M., Brown, M. A., McCarthy, M. I., Yang, J., 2012. Five years of gwas discovery. The American Journal of Human Genetics 90 (1), 7–24.

Witte, J. S., Visscher, P. M., Wray, N. R., 2014. The contribution of genetic variants to disease depends on the ruler. Nature Reviews Genetics 15 (11), 765–776.

Wray, N. R., Gottesman, I. I., 2012. Using summary data from the danish national registers to estimate heritabilities for schizophrenia, bipolar disorder, and major depressive disorder. Frontiers in genetics 3.

Yang, J., Benyamin, B., McEvoy, B. P., Gordon, S., Henders, A. K., Nyholt, D. R., Madden, P. A., Heath, A. C., Martin, N. G., Montgomery, G. W., Goddard, M. E., Visscher, P. M., Jul 2010. Common SNPs explain a large proportion of the heritability for human height. Nat. Genet. 42 (7), 565–569.

Yang, J., Lee, S. H., Goddard, M. E., Visscher, P. M., 2011a. Gcta: a tool for genome-wide complex trait analysis. The American Journal of Human Genetics 88 (1), 76–82.

Yang, J., Weedon, M. N., Purcell, S., Lettre, G., Estrada, K., Willer, C. J., Smith, A. V., Ingelsson, E., O’Connell, J. R., Mangino, M., et al., 2011b. Genomic inflation factors under polygenic inheritance. European Journal of Human Genetics 19 (7), 807–812.

Zhou, X., Carbonetto, P., Stephens, M., 2013. Polygenic modeling with bayesian sparse linear mixed models. PLoS genetics 9 (2), e1003264.

Zhou, X., Stephens, M., 2014. Efficient multivariate linear mixed model algorithms for genome-wide association studies. Nature methods 11 (4), 407–409.

Zöllner, S., Pritchard, J. K., 2007. Overcoming the winner2019s curse: estimating penetrance parameters from case-control data. The American Journal of Human Genetics 80 (4), 605–615.

